# Using social information improves individual foraging efficiency in sheep

**DOI:** 10.1101/2023.08.05.552097

**Authors:** Vartan E Vartparonian, Stephan T Leu

## Abstract

Knowledge about the environment is fundamentally important to move, find resources and forage efficiently. This information can either be acquired through individual exploration (personal information) or from other group members (social information). We experimentally assessed the use of social information and its influence on foraging efficiency in sheep, *Ovis aries*. Naïve individuals paired with an informed partner that knew the food patch location, found the patch significantly faster compared to naïve individuals paired with another naïve individual. Similarly, they spent a significantly lower proportion of time exploring areas away from the food patch. We further found that the previous outcome of using social information (success = access to feed vs failure = no access to feed) had no impact and sheep continued to use social information in subsequent foraging trials and foraged similarly efficient. Our results suggest, naïve sheep that are unfamiliar with resource locations, forage more efficiently when informed individuals are present compared to when all individuals are naïve. If informed individuals play a similar role in larger groups, new management practices could be developed to improve foraging efficiency when sheep are moved to new paddocks or in paddocks with heterogenous and dynamic resource distribution.

## 1. Introduction

Environmental conditions are often rapidly changing which can result in a variable and uncertain food resource distribution (Nichols et al., 2011; Baskin et al., 2014). As a consequence, animals must modulate their foraging behaviour and learn where and when it is most effective to forage. Animals can learn about their surrounding environment by collecting personal information, through exploration and interacting with the environment. For instance, individuals collect personal information regarding the distribution of resources by sampling different food patches while foraging (Valone and Templeton, 2002; Kendal et al., 2005). This sampling technique provides reliable information but requires time and energy (Bonnie and Earley, 2007). Animals can also collect social information, through observing other individuals interact with the environment (Dall et al., 2005), and for instance, obtain social information about resource distribution (Bonnie and Earley, 2007; Valone, 2007; Duboscq et al., 2016). Following informed individuals to locate food resources is often faster than finding it themselves (Held et al., 2000; Danchin et al., 2004; Bonnie et al., 2007). Hence, using social information, compared to personal information, is typically less costly (Bonnie and Earley, 2007), but may not always be reliable.

The use of social information has been shown in several wildlife taxa (birds, fish, mammals; Templeton and Giraldeau, 1996; Reader et al., 2003), as well as livestock species (goats, cattle, and pigs; Held et al., 2000; Shrader et al., 2007; Costa et al., 2016). For example, tamarins can learn the location of a food patch by monitoring the behaviour of other conspecifics (Bicca-Marques and Garber, 2005) and young unmated dairy cattle begin their grazing activity quicker in the presence of older and more experienced group members (Costa et al., 2016). However, our understanding of social information use in sheep and whether it is beneficial is limited. Nevertheless, we do know that sheep collect personal information when foraging, allowing them to gain knowledge about the resource distribution within a paddock (Edwards et al., 1996; Dumont and Petit, 1998). Sheep use spatial cognition, memory, and landmarks to remember and distinguish between different food patches, and then move back to those patches during subsequent foraging bouts (Edwards et al., 1996; Dumont and Petit, 1998). However, when the distribution of resources becomes more random, and less predictable, repeated use decreases, because it requires more time to locate these random patches (Edwards et al., 1996). In many habitats, resource distribution is patchy, and changes with season. Furthermore, 10% of the Australian sheep population lives in arid regions with variable distribution of resources (Department of Agriculture and Water Resources, 2006). Therefore, the use of social information while foraging could be advantageous for sheep and improve their foraging efficiency.

Sheep are often moved between paddocks. For younger sheep, this includes paddocks they have not experienced before. Lambs that experience unfamiliar food options, have been shown to have reduced foraging efficiency and feed intake compared to lambs that had previous exposure to the feed (Flores et al., 1989). Even if sheep have been in a paddock before, the resource distribution may have changed over time and with seasons, making their previously collected information unreliable (Edwards et al., 1996).

Using social information can improve foraging efficiency (Templeton and Giraldeau, 1996; Valone and Templeton, 2002). However, when social information is unreliable, the use of personal information is preferred (van Bergen et al., 2004; Kendal et al., 2005). We hypothesised that (i) sheep use social information in a foraging context, and that this improves their foraging efficiency; and (ii) if sheep were unsuccessful using social information, they reduce the use of social information during subsequent foraging. Our findings will provide deep insight into the influence of social behaviour and in particular, the use of social information on foraging decisions and foraging efficiency. If naïve sheep forage more efficiently in the presence of informed conspecifics, then new management practices, such as adding informed sheep to naïve mobs, could be developed to utilise this relationship.

## 2. Materials & Methods

### 2.1. Study site and animals

This study was carried out at The University of Adelaide, Roseworthy Campus, South Australia, in the Austral winter of 2022. We observed 80 female sheep (Merino, Border Leicester crosses) of similar age (1 to 2 years). We randomly divided the 80 sheep into five groups of 16 individuals, where each group was housed in a separate large paddock. Each group was then observed for a five-day period. At the end of the observation, sheep were released back into their original paddock. First, we selected eight of the 16 individuals from each group and used them during the first phase (training phase). These eight individuals were separated into four pairs, each housed in a separate yard. The two individuals of each pair were housed together between 17 to 20 hours before their first trial and had similar familiarity with each other. The other eight animals remained in their large paddock and were used during the second phase (testing phase).

### 2.2. Arena

We conducted the behavioural observations in an arena 65 × 38 metres (0.25 ha; Figure 1). The arena was surrounded by privacy netting so the tested sheep could not see any animals in neighbouring paddocks, minimizing the potential effect of external factors. A midline fence, 25 metres in length, separated the end zone into two sections (bucket with food, bucket without food). This division aims to stimulate the decision making of sheep when selecting a side during foraging and allows to measure the proportion of time spent on the incorrect side (*Error Proportion*), i.e., where the feed bucket was empty (Lee et al. 2006). Cameras were placed at opposite ends of the arena, approximately 6 metres outside the arena. The cameras filmed all trials from an elevated viewpoint (approx. 6m high). Directly adjacent to the testing arena was a holding area, which we used to allow the two sheep to settle for up to five minutes after being moved to the testing arena, and prior to the start of the observation. During the five minutes sheep were allowed to walk unassisted into the testing area through the open gate, which started the observation period. If sheep didn’t do that, they were gently moved into the testing arena after five minutes.

**Figure 1.**
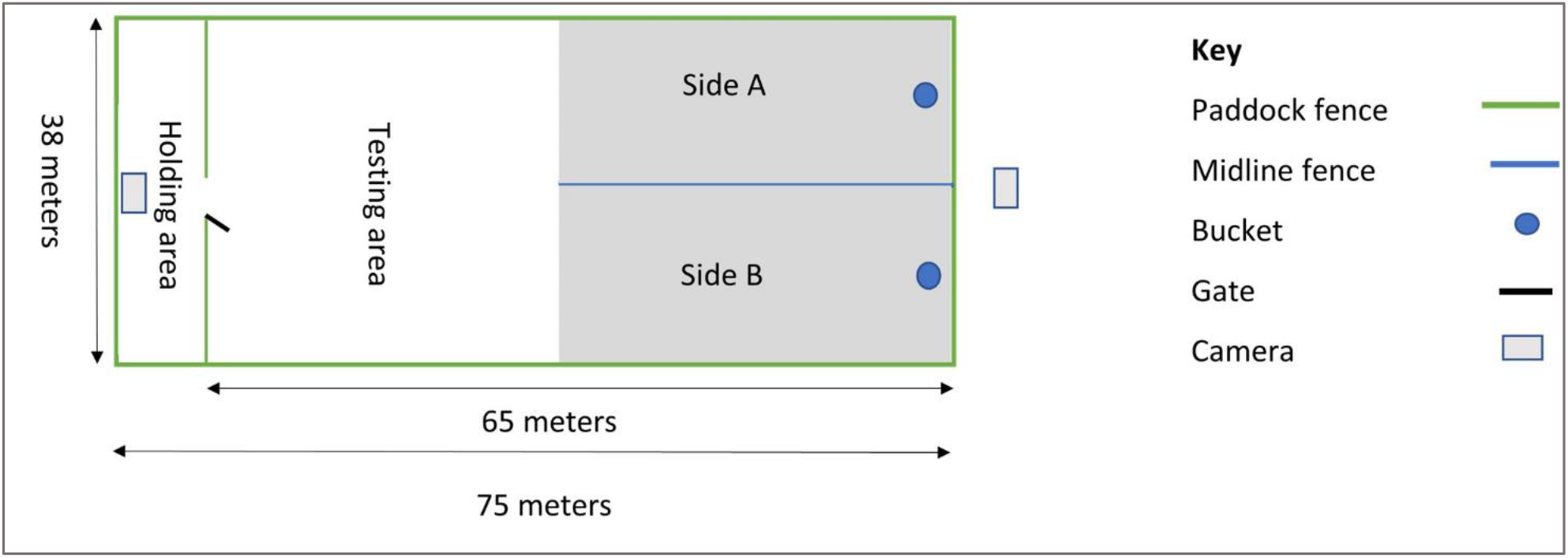
Diagram of arena used for testing. The endzone, grey shaded area, was used to calculate the ‘Error Proportion’.

### 2.3. Experimental design

#### 2.3.1. Phase 1, training phase

On day 1 and 2, the pairs were trialled twice each day to learn the location of the feed bucket. We placed one bucket on each side of the arena (side a and b; Figure 1). Only one bucket was filled with hay, the other was empty. Half of the pairs had the bucket filled with hay on the left side while the other half had it on the other side to account for any potential side differences. Pairs that did not locate the feed bucket within the 45-minute trial period were led to the feed bucket and allowed to feed from it to assist learning the location. This was required in thirteen of the twenty trials (65%) during trial 1 and decreased to seven of the twenty trials (35%) during trial 4. The very first trial of each pair, when both sheep were completely naïve to the feed location, acted as the control. On day 3, trial 5 and 6, we assessed whether sheep had learned the feed location, a prerequisite for the sheep to be used during the next observation. We used the following criterion to classify sheep as informed: During at least one of the two trials (5 or 6), the sheep needed to locate the feed bucket in less than 15 minutes and without entering the incorrect side. Only the sheep that was leading and found the feed bucket first, was selected for the following observations. This ensured that this individual knew about the location of the feed bucket itself and did not simply follow its partner. Each of the four selected informed sheep was then housed with two of the eight naïve sheep. The other four individuals that were not selected were removed from the study and released back to the paddock.

#### 2.3.2 Phase 2, naïve sheep paired with informed partner

In this phase, we quantified the foraging efficiency of a naïve sheep paired with an informed individual. On day 4, the informed sheep were paired and trialled once with each of the two naïve individuals they were housed with. Half of the pairs had access to the feed bucket, making it successful. The other half were unable to access the feed bucket filled with hay as it was covered with wire mesh, making it unsuccessful. On day 5, the same trials were run again, but feed was always accessible. Day 4 and 5 together, allowed us to determine whether continued use of social information was dependent on previously using it successfully and whether it influenced foraging efficiency (Figure 2).

**Figure 2.**
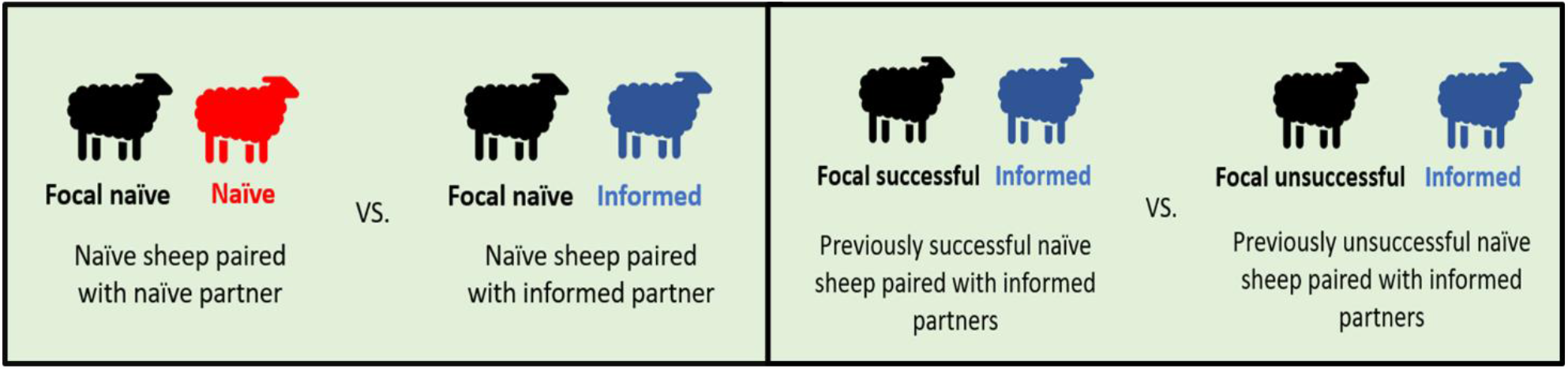
Study design. Data from focal (black) naïve sheep paired with a naïve (red) partner or informed (blue) partner were used to address whether the presence of an informed individual influences the foraging efficiency of a naïve individual. Comparing data from focal successful and unsuccessful sheep were used to assess whether the subsequent foraging efficiency and social information was influenced by the previous outcome (successful vs unsuccessful). Data collected from black ‘focal’ sheep were used for data analyses.

### 2.4. Video and GPS tracking of animals

Each sheep was equipped with a coloured blanket attached over its back, and with a Global Positioning System (GPS) collar. The blanket (red or grey) allowed us to individually identify each sheep on the video recording. The collars included a GPS unit (i-gotU GT-120, by MobileAction, with a larger battery CE04381 by Core Electronics). We used the GPS coordinates to calculate distances between both sheep (details below). At the end of each day, we replaced the GPS collars with fully charged ones to record the trials of the following day. The total weight of each collar was approximately 700 grams, or 1.1% of the mean live weight of our sheep (61.3 ± SE 0.8 kg). All sheep were handled using procedures formally approved by The University of Adelaide Animal Ethics Committee in compliance with the Australian code for the care and use of animals for scientific purposes (approval number S-2022-024).

### 2.5. Behavioural measures

We used the duration to find the feed bucket as a measure of foraging efficiency. For each individual we determined the *Total Time* as the time period from entering the testing area until the feed bucket was located. If an individual did not find the bucket within the observation period of 45 minutes, the trial was terminated, and its *Total Time* recorded as 45 minutes. For each pair, we also recorded the *Time Difference* between the *Total Time* of sheep 1 and sheep 2. As a complementary measure of foraging efficiency, we determined the *Error Proportion*, which we calculated as the proportion of time spent on the incorrect side of the endzone relative to the time spent in the endzone (Figure 1). To determine whether social information, that is the presence of an informed conspecific, influenced foraging efficiency, we compared the *Total Time* and *Error Proportion* of naïve sheep when paired with another naïve sheep (day 1, trial 1) versus when paired with an informed sheep (day 4). Similarly, we compared these times of the final trial (day 5) between previously successful and unsuccessful sheep. Following the unsuccessful foraging bout, behavioural changes may be subtle, and sheep still paying attention to the informed individual but follow them less closely. That is the inter-individual distance may be larger. Therefore, we used the GPS data to calculate the *Inter-individual Distance* between pair partners during the final trial (day 5) and compared it between sheep that had been successful versus unsuccessful in the earlier trial (day 4).

### 2.6. Statistical Analysis

All statistical tests were conducted using IBM® SPSS® Statistics 27. The data were not normally distributed for any of the treatment groups (Shapiro-Wilk test, P-value <0.05). Therefore, we used a Mann-Whitney U-Test to compare measurements between treatment groups.

## 3. Results

### 3.1. The effect of an informed partner on foraging efficiency

We only used the data when sheep had access to the feed in the bucket, so the trial conditions were identical for the different pairings. Naïve sheep that were paired with an informed partner foraged more efficiently than sheep paired with a naïve partner. These individuals found the feed bucket (*Total Time*) significantly faster (Mann Whitney U Test, U=87, p=0.003, N=39, Figure 3). We excluded one observation from the analysis (resulting in N=39 observations) as one naïve individual found the feed bucket before its informed partner. The majority of naïve individuals did not locate the feed bucket within the allocated observation time when paired with a naïve partner, indicated by a median *Total Time* of 45 minutes. Consistent with the shorter *Total Time*, when paired with an informed individual, naïve sheep spent a smaller proportion of time on the incorrect side of the endzone (*Error Proportion*, Mann Whitney U Test, U=75, p=0.028, N=33) compared to those paired with a naïve partner (Figure 4). Some pairs did not enter the endzone during the trial, and were removed from the analysis, reflected by the lower sample size.

**Figure 3.**
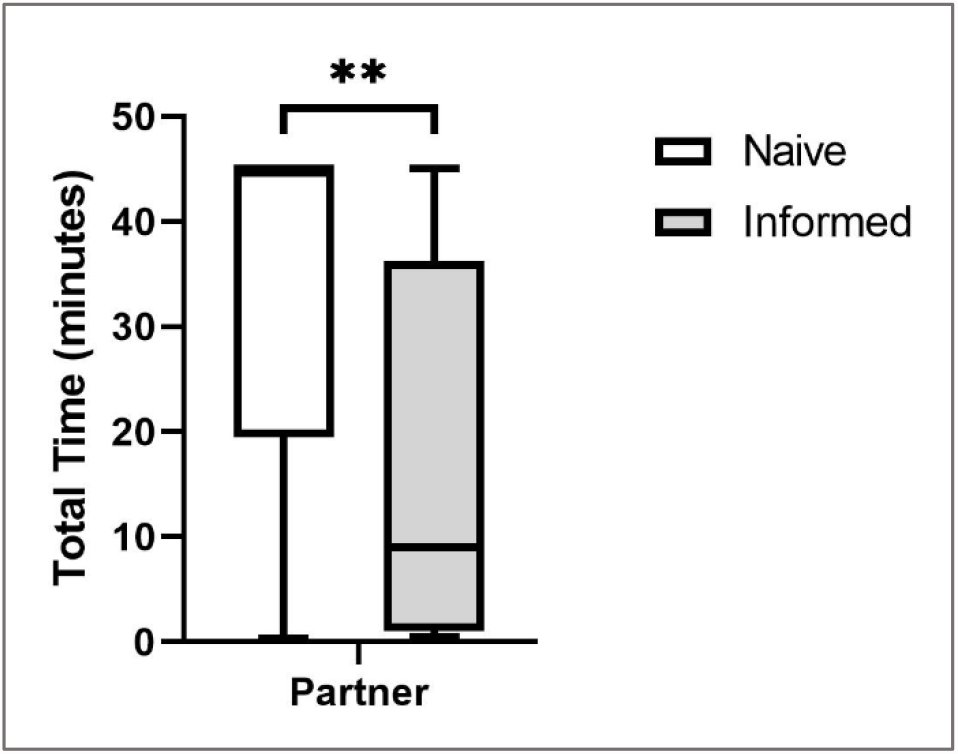
The Total Time to find the feed bucket was significantly faster when naïve individuals were paired with an informed partner (grey bar) compared to being paired with a naïve partner (white bar). ** = significant difference (p-value <0.01).

**Figure 4.**
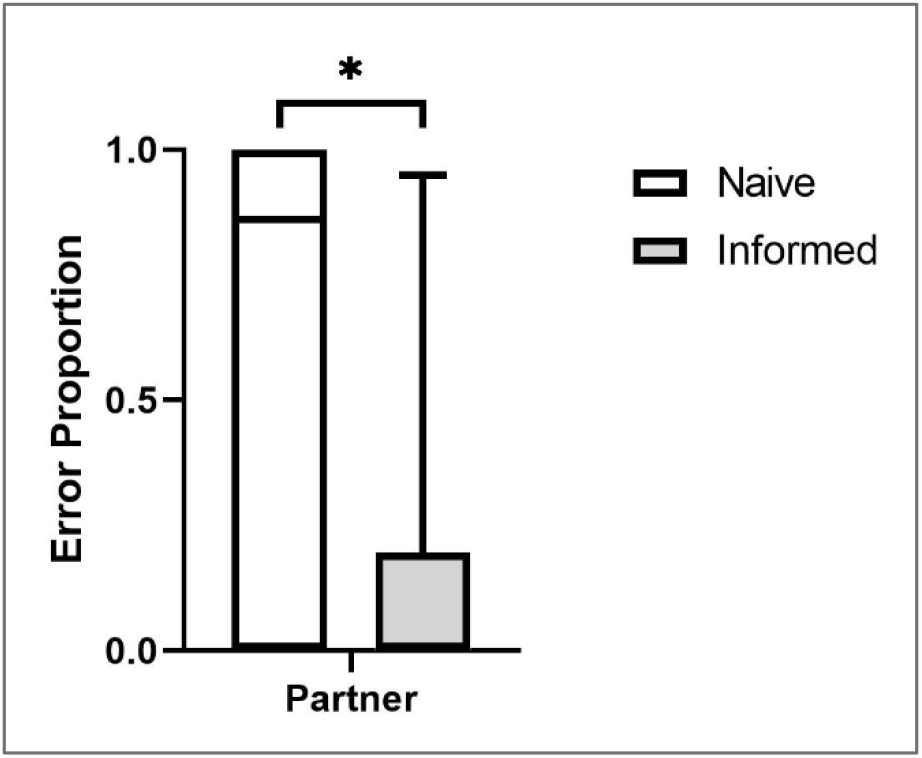
Error Proportion for naïve sheep was significantly lower when paired with an informed partner (grey bar) compared to being paired with a naïve partner (white bar). * = significant difference (p-value <0.05).

### 3.2. The immediate effect of the unsuccessful use of social information

The *Time Difference* between pair partners finding the feed bucket was similar irrespective of whether the previous foraging trial was successful (access) or unsuccessful (no access to feed; Mann Whitney U Test, U=58, p=0.072, N=28; Figure 5). When neither of the pair partners found the bucket (N=11), we removed the data from the analysis. Additionally, we removed one outlier data point, because its value (8.42 min) was more extreme than Q3+1.5(IQR) = 1.16 min (Bakker and Wicherts, 2014). Finally, the inter-individual spatial distance between pair partners did not differ significantly between the group that had access to feed in the previous trial and the group that did not have access to feed (Mann Whitney U Test, U=82, p=0.496, N=28, Figure 6).

**Figure 5.**
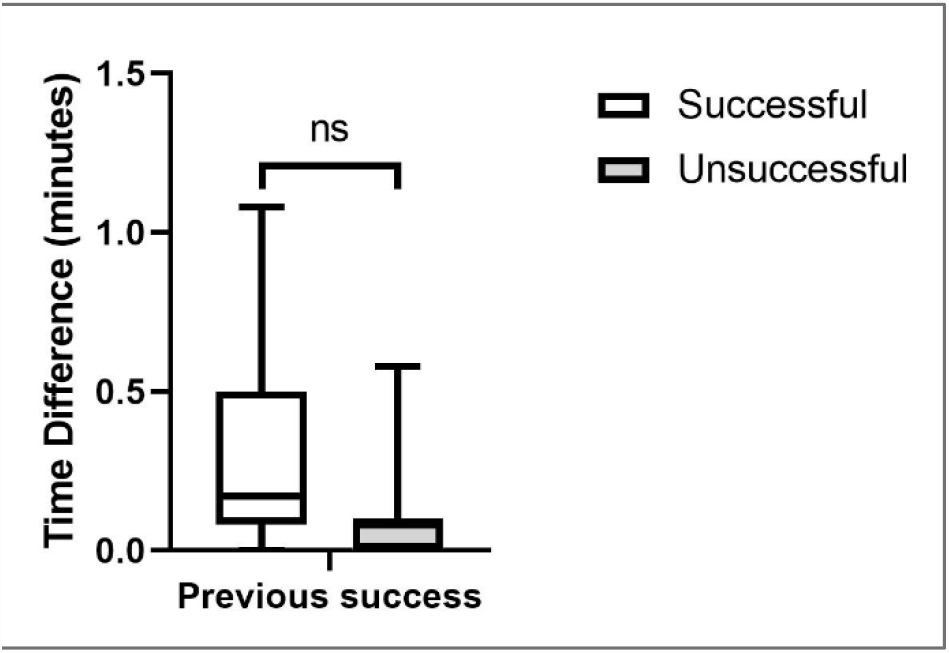
The Time Difference between pair partners finding the feed bucket did not depend on whether the previous foraging trial was successful (access to feed) or unsuccessful (no access to feed). There was no significant difference in Time Difference between treatment groups (ns = not significant).

**Figure 6.**
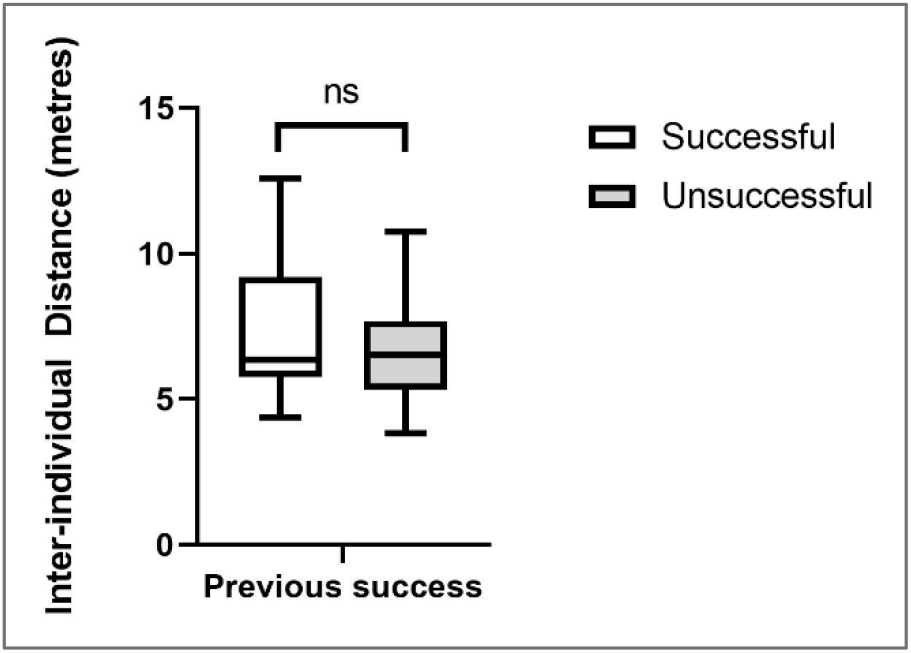
The Inter-individual Distance between pair partners was not significantly different between sheep that were previously successful (access to feed) or unsuccessful (no access to feed; ns = not significant).

### 3.3. The effect of previous success on subsequent foraging efficiency

Previous success had no immediate effect on the foraging efficiency of sheep during the subsequent foraging trial. There was no significant difference in *Total Time* (Mann Whitney U test, U=135.5, p=0.284, N=37), nor *Error Proportion* (Mann Whitney U test, U=100.5, p=0.446, N=31) between treatment groups. We excluded some data from the *Total Time* analysis, because in two cases, the naive sheep reached the feed bucket before their informed partner, and in one case the incorrect naïve sheep was trialled. The Error Proportion analysis only had 31 data points as nine pairs did not enter the endzone throughout the entire trial.

## 4. Discussion

Foraging efficiency was improved when naïve sheep were paired with an informed partner compared to being paired with another naïve individual. They found the feed bucket significantly faster (shorter *Total Time*) and the proportion of time spent on the wrong side was lower (*Error Proportion*). Social information transmission and social learning could explain this finding. Animals can acquire information about their environment by observing conspecifics (Giraldeau et al., 2002; Dall et al., 2005) and this can influence foraging efficiency in both wild and domestic species (Bonnie and Earley, 2007; Shrader et al., 2007; Jaeggi et al., 2010; Costa et al., 2016). For example, Shrader et al. (2007) observed that goats selected a tray with the largest amount of food instead of two other trays with less food when they were able to see a conspecific feeding from those same trays beforehand. And Held et al. (2000) observed that naïve pigs inspected fewer buckets to locate the one with food, when they followed an informed pig. In sheep, movement can be a cue to conspecifics and has been shown to recruit other individuals to follow (Bonnie et al., 2007). For example, Pillot et al. (2010) showed that one individual that was trained to move towards a sound cue, was followed by other and untrained group members. A similar mechanism could drive our findings. Due to its knowledge where the feed bucket was, the informed partner could have walked straight towards the bucket. This directional movement towards a location could provide a cue to the naïve individual to follow, which would reduce the *Total Time* and *Error Proportion*.

An alternative, but not mutually exclusive explanation for the increased foraging efficiency is the gregariousness of sheep and their propensity to maintain social cohesions (Dumont and Boissy, 2000; Dumont and Hill, 2001; Boissy and Dumont, 2002). For instance, Dumont and Boissy (2000) showed that sheep walk away from conspecifics to a preferred foraging patch if it was within a 15-metre distance but did not if it was 50 metres away. Site familiarity further modulates social cohesion (Warren and Mysterud, 1993; Scott et al., 1996; Rook et al., 2005). For instance, lambs are more likely to move away from their social group to feed on their preferred food items when they were familiar with the paddock (Scott et al., 1996). In unfamiliar paddocks the social group stayed together (Scott et al., 1996). In our study, the informed sheep were able to familiarise themselves with the arena during the six trials of the training phase. Hence, the informed sheep may have prioritised accessing the feed over maintaining social cohesion and quickly moved to the feed bucket after entering the testing arena. In contrast, the naïve partners were unfamiliar with the testing arena during their first trial (day 4). Therefore, they may have prioritised social cohesion with the informed individual (Scott et al., 1996). This would suggest that the observed foraging efficiency of naïve sheep was improved due to their sociability and not through the transmission of social information about the feed location. Nevertheless, irrespective of the mechanistic driver, both possible explanations show the importance of informed conspecifics to improve foraging when resource distribution is heterogeneous.

The use of social information may be modulated if this strategy was previously unsuccessful. However, we did not find any empirical evidence that sheep used personal information more strongly after social information use was unsuccessful in the previous trial. Inter-individual spatial distance, as well as the time difference between pair partners arriving at the feed location were not significantly different irrespective of whether the previous trial was successful or not. The spatial and time differences are related to each other, because sheep that are further away from their partner take longer to reach the bucket. Hence, both results are consistent with each other.

Another possible explanation for a lack of effect is that one unsuccessful trial is not enough to change the foraging strategy and subsequent use of social information. This notion is further supported by our finding that foraging efficiency (*Total Time* and *Error Proportion*) was not affected by the success of the previous foraging bout. Furthermore, after only one trial, the naïve sheep are still relatively unfamiliar with the environment. Hence, they may use the strategy ‘copy when uncertain’, where animals use social learning (social information) when they have no relevant prior knowledge. Hence, they may adopt the foraging behaviour of their conspecifics when in an unfamiliar environment (Laland, 2004). However, it is conceivable, if following an informed individual repeatedly results in unsuccessful foraging, sheep may ultimately determine the social information as unreliable, and prioritise personal information instead (Kendal, 2004). For instance, van Bergen et al. (2004) have shown this in sticklebacks, which use personal information when it is reliable instead of social information. However, when personal information is unreliable (56% success rate), stickleback use both type of information (van Bergen et al., 2004).

As before, sociability could drive the lack of effect, if naïve sheep simply follow informed sheep in order to maintain social cohesion irrespective of foraging. Boissy and Dumont (2002) observed that when in pairs, sheep remained close to each other instead of approaching a preferred foraging patch. When more than two sheep were present in the group, they were more likely to exploit preferred foraging patches and move away from the group, but mostly in small subgroups and not alone. Hence, in our study, the lack of alternative social partners and its social propensity may have influenced the naïve sheep to stay close to the informed sheep irrespective of the success or failure of the previous foraging bout.

## 5. Conclusion

Our study showed that the presence of an informed individual improves the foraging efficiency of naïve individuals. Two mechanisms could explain our findings. Naïve sheep were driven by their social propensity and therefore indirectly obtained information about the new environment by remaining close to their informed conspecifics. Alternatively, naïve sheep actively used social information and obtained information about the resource distribution in a new environment by observing the behaviour of their informed partner. Irrespective of the mechanism, there was a clear benefit on foraging efficiency of the presence of an informed individual. However, we only showed this in a pair context. Social groups in sheep are typically larger and can reach up to 150 members in wild sheep (Shackleton and Shank, 1984) and many hundreds on farms. However, sheep have a dynamic fission-fusion social system and during periods of activity sheep temporarily split into much smaller sub-groups (Dela Libera et al., 2023). Nevertheless, further investigation of the use of social information in larger groups would provide further insight. It would be interesting and important to identify the ideal ratio of informed to naïve individuals that increases the effectiveness of social foraging. Aplin et al. (2015) showed that a very small number of informed individuals can affect large populations and even introduce new foraging behaviours. If our results upscale to larger groups, using informed individuals to improve foraging could be an interesting approach for the sheep industry to develop new management practices. For example, when moving sheep to a new paddock, adding individuals that have had previous exposure to this site may improve the foraging efficiency, and ultimately weight gain of newcomers.

## 6. Acknowledgements

We would like to thank the staff of the Roseworthy Campus Farm (Martindale Holdings) for logistical support during the project. Vartan Vartparonian was supported by the Davies Livestock Research Centre at the University of Adelaide. This work and Stephan Leu were supported by an Australian Research Council DECRA Fellowship [DE170101132].

